# A multivalent mRNA-lipid nanoparticle vaccine containing eight hemagglutinin antigens elicited broad neutralizing antibody responses and protected against influenza A virus challenge in swine

**DOI:** 10.64898/2026.06.08.730105

**Authors:** Giovana Ciacci Zanella, Marissa L. Vincent, Leo Flores, Ethan K. Aljets, Rodrigo C. Paiva, Alexey Markin, Blake Inderski, Garima Dwivedi, Drew Weissman, Meghan Wymore Brand, Jefferson J. S. Santos, Scott E. Hensley, Tavis K. Anderson, Phillip C. Gauger, Amy L. Baker

## Abstract

The diversity within H1 and H3 subtype influenza A viruses (IAV) in swine prevents effective vaccine control approaches with inactivated whole-virus vaccines. We addressed the challenge of controlling co-circulating hemagglutinin (HA) clades of swine IAV with the development of a multivalent mRNA-lipid nanoparticle (LNP) vaccine expressing 8 HA proteins to maximize genetic coverage. We applied a computational approach to select eight HA genes that represented 95% of the observed IAV detected in the United States between 2022 and 2025. Piglets were vaccinated and boosted intramuscularly with either individual HA mRNA-LNP or an 8-HA multivalent mRNA-LNP. Serum was collected to evaluate systemic antibody levels. Twenty-one days post-boost, pigs were challenged with a field relevant H1 1A.3.3.3-c3 IAV strain. The 8-HA multivalent mRNA-LNP vaccine induced neutralizing antibodies against all eight antigens and vaccinees were protected against lung lesions, with lesion scores similar to non-challenged animals. Homologous monovalent vaccination significantly reduced IAV detection in nasal secretions and in the lungs. Heterologous monovalent vaccination was not cross-protective but did not induce vaccine-associated enhanced respiratory disease. We provide evidence that monovalent and multivalent mRNA-LNP influenza vaccines elicited neutralizing antibody responses in pigs and protected against viral challenge. The versatility and capacity for rapidly updating the mRNA-LNP vaccine platform make it an appealing tool to improve animal health and minimize the circulation and diversity of IAV in swine.

**Importance:** Influenza A virus is an important respiratory pathogen in swine, and zoonotic transmission of swine strains to humans remains a public health risk. Control strategies against IAV in swine herds rely heavily on biosecurity measures and vaccination. However, the antigenic diversity of IAV circulating in swine challenges current vaccination programs, and there is a need for broadly protective vaccines or platforms that can rapidly update components to reflect circulating diversity. mRNA-LNP vaccines have emerged as promising vaccine platforms, offering simultaneous delivery of multiple antigens, rapid development, scalable manufacturing, and potent immunogenicity. In this study, we assessed the immunogenicity and protective capacity of monovalent and multivalent mRNA-LNP vaccines encoding eight representative IAV HA antigens. To our knowledge, this is the first study to objectively select multiple representative endemic swine IAV strains by quantifying genetic diversity within the phylogeny and to apply this selection to rationally design and evaluate a multivalent HA mRNA-based influenza vaccine in the swine model.

## Introduction

Respiratory disease caused by influenza A viruses (IAV) of the H1N1, H3N2, or H1N2 subtypes is a significant health and economic burden for the swine industry [1]. In addition to clinical complications and production impacts, the circulation of IAV in swine represents a public health concern [2]. Pigs have played a significant role in IAV ecology and evolution by serving as “mixing vessels” in which avian, human, and swine strains may exchange gene segments in a process known as reassortment [3]. Reassortment events can create novel IAV genotypes that may have altered phenotypes that can cross species barriers and potentially lead to pandemics [4]. In the United States, repeated interspecies spillovers, followed by evolution in pigs has resulted in antigenic drift and significant strain variation between regions and within swine production systems [5]. The HA and NA genes of IAVs regularly detected in swine in the United States (U.S.) are classified into more than 20 distinct HA and 11 NA clades by phylogenetic analysis [6, 7]. This diversity, specifically of the surface glycoproteins, poses a significant barrier to controlling infection and the development of broadly protective vaccines [8].

Control strategies against IAV in US swine rely heavily on biosecurity measures and vaccination. The majority of IAV vaccines utilized are designed to induce systemic HA-specific antibodies and are most effective when the HA of the vaccine strains closely matches that of circulating wild-type strains [5]. Since a significant challenge to effective commercial vaccines is the concurrent circulation of multiple IAV genetic clades within the same region, and there is no centralized selection of annual vaccine strains for the swine industry, customized autogenous and prescription vaccines are commonly used by U.S swine producers [9]. Custom vaccine formulations are informed by production systems or regional genomic surveillance and are more likely to be well-matched. HA-specific antibodies that are well matched and targeted to the receptor-binding site can block viral attachment to host cell receptors and neutralize viral infectivity, thereby effectively reducing viral load, clinical disease, and transmission [10, 11].

The two IAV vaccine platforms licensed and available for use in the swine industry are whole inactivated virus (WIV) vaccines and non-replicating alphavirus RNA particle (RP) vaccines. WIVs were historically the most widely used platform for controlling swine IAV [9]. WIV reduce clinical disease and lung lesions and provide partial reduction of shedding when the vaccine antigen and challenge viruses are antigenically similar [12]. However, the substantial genetic and antigenic diversity of endemic swine IAV limits the ability of these vaccines to provide the breadth of protection needed. WIV vaccines are susceptible to interference from maternally derived antibodies (MDA) [13], have limited ability to induce robust cellular immune responses [8], and were associated with exacerbated clinical signs and vaccine-associated enhanced respiratory disease (VAERD) when mismatched [12, 14]. In addition, for a highly diverse and rapidly evolving pathogen such as IAV, a major challenge of current vaccine approaches is the ability to rapidly update and produce vaccines in response to emerging antigenically novel strains [15]. Because WIV vaccines require virus isolation and culture in eggs or cells in laboratories with appropriate biosafety levels, this results in a longer production timeline and limits the speed at which updated vaccines can be produced and distributed at a large scale [16]. Alphavirus-derived RP technology offers several advantages for controlling IAV in comparison to WIV vaccine platforms in swine. By using replication-defective viral particles to deliver antigen sequences, these vaccines evoke strong intracellular antigen expression and robust humoral and cellular immune responses without causing VAERD [17–19]. This technology also allows for the inclusion of multiple IAV antigens and has the benefit of being based on the HA or NA sequences, eliminating the need for the virus isolation step and therefore shortening the production timeline [18, 20].

Although commercial, autogenous WIV and prescription RP vaccines are available and used in over 80% of U.S breeding herds, 74.2% of surveyed veterinarians agree on the need of new or improved vaccine platforms for the swine industry [21]. This is likely due to the IAV challenges faced in the field and the limitations of current vaccines. There is a demand for additional improved IAV vaccination strategies in commercial swine that induce the broadest possible immune response and overcome interference from MDA, while still providing adequate mucosal and cell-mediated immune responses following heterologous infection without causing VAERD. Responding to these demands by using conventional vaccine approaches may be difficult, but they may be achievable by investigating alternative nucleic acid-based vaccine platforms.

Messenger RNA (mRNA) vaccines encapsulated in lipid nanoparticles (LNPs) have emerged as promising alternatives that offer simultaneous delivery of multiple antigens, rapid development, scalable manufacturing, and potent immunogenicity [16]. Antigens from 20 distinct HA genes were simultaneously delivered via mRNA-LNPs to mice and ferrets and were efficient at eliciting broad humoral and cellular immunity [22]. In this study, our objective was to extend this approach to the swine model based on epidemiologically relevant HA clades selected through an algorithm that identifies genetically representative genes [23]. To assess the immunogenicity and protective capacity of mRNA-LNP vaccines, piglets were vaccinated and boosted intramuscularly with either an individual HA mRNA-LNP or an 8-HA multivalent mRNA-LNP, and a subset of the groups were challenged. Our work provides evidence that mRNA-LNP vaccines elicited expression of multiple HA targets and induced humoral immune responses to each HA in pigs in a clade specific manner. This vaccine approach protected against clinical disease and reduced viral replication and shedding following challenge. To our knowledge, this is the first study to objectively select multiple representative endemic swine IAV strains by quantifying genetic diversity within the phylogeny and to apply this selection to a multivalent HA mRNA-LNP influenza vaccine in the swine model.

## Results

### Piglets vaccinated with the multivalent mRNA-LNPs produced HI antibody titers against all 8 encoded HAs

Eight HA vaccine antigens were selected from the most frequently detected HA clades within genomic surveillance; the selected antigens provided genetic coverage of 63.61% of H1 (n=2,674) and 71.25% of H3 (n=1,419) HA genes detected between 2022-2025 in the U.S. (Figure 1). The sequences of these antigens were then used to prepare eight HA-encoding nucleoside-modified mRNAs encapsulated in LNPs. IAV antibody naïve piglets (N=2) were vaccinated and boosted intramuscularly with a 50 µg dose of each individual HA mRNA vaccine (monovalent) to verify the immunogenicity of each vaccine component. In addition, another group of piglets (N=10) was vaccinated with 50 µg of the multivalent IAV mRNA-LNPs that included 6.25 µg of each of the 8 HA mRNA-LNPs to evaluate simultaneous antibody responses against all eight encoded HAs. Serum was collected at multiple time points to assess vaccine response by hemagglutination inhibition (HI) assays (Figure 2). All eight monovalent high-dose HA mRNA-LNP vaccines induced antibodies that recognized homologous wild type viruses in a clade-specific manner (Table 2 and 3). Piglets vaccinated with the monovalent mRNA-LNP vaccines had antibody titers at or above 1:40 by 14 days post-vaccination (DPV) that increased significantly 14 days after the boost dose. The piglets that received the multivalent 8-HA mRNA-LNP vaccine produced antibodies that reacted to all eight encoded HA antigens following the boost administration, albeit at lower levels relative to piglets that received 50 µg of each monovalent HA mRNA-LNP vaccine.

**Figure 1.**
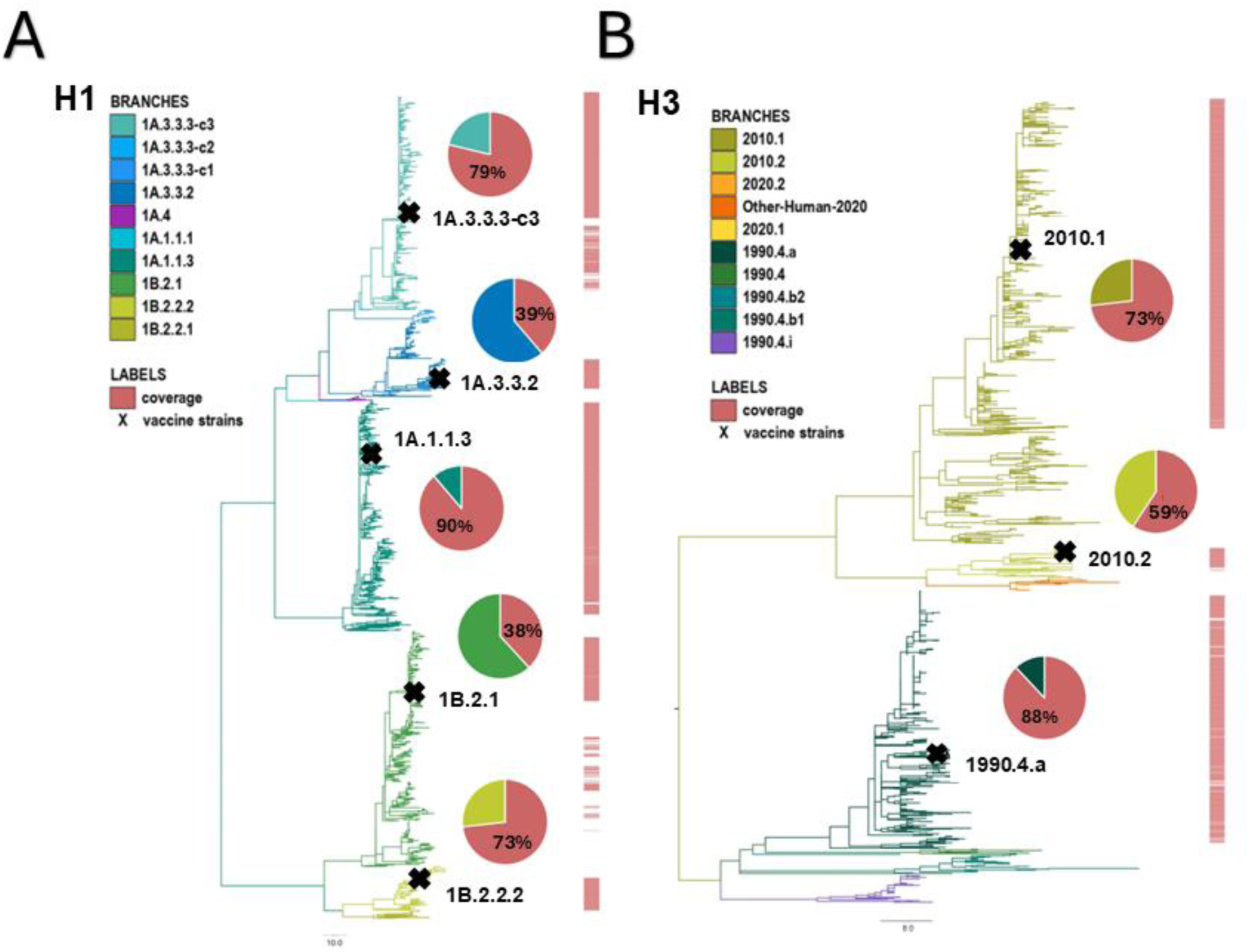
Phylogenetic trees of H1 (A) and H3 (B) collected from the U.S. between 2022 and 2025, with branches rescaled by protein HA1 alignment. For each vaccine strain (X), clade coverage was determined at a 16 amino acid (∼5%) similarity threshold using PARNAS and is displayed in the pie chart. The 8 selected HA genes provided coverage of 63.61% of H1 (n=2,674) and 71.25% of H3 (n=1,419) genes.

**Figure 2.**
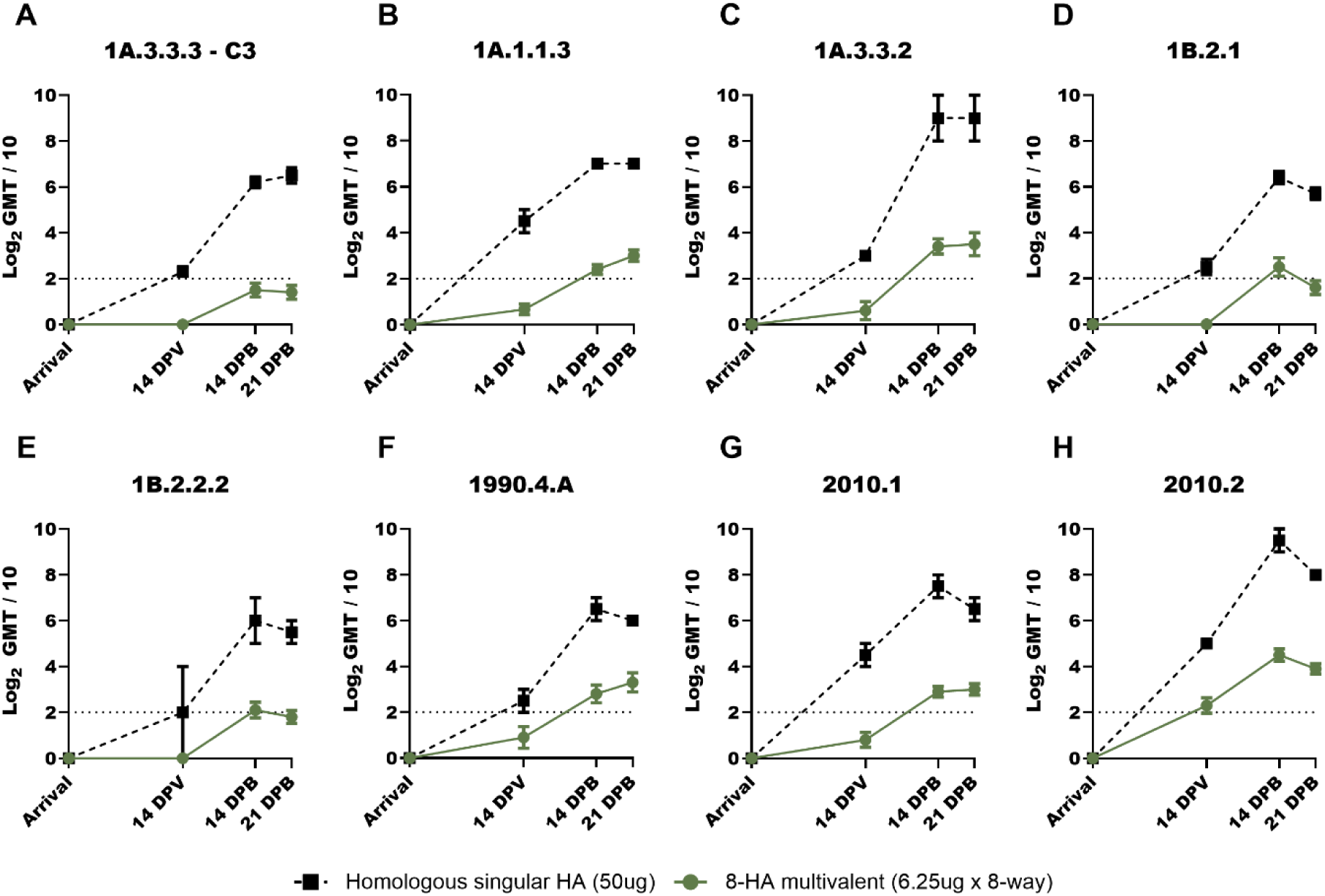
Hemagglutination inhibition (HI) antibody responses against individual HA antigens following vaccination. Log2-transformed geometric mean HI titers against (A) 1A.3.3.3-c3, (B) 1A.1.1.3, (C) 1A.3.3.2, (D) 1B.2.1, (E) 1B.2.2.2, (F) 1990.4a, (G) 2010.1 and (H) 2010.2 are shown for pigs vaccinated with homologous singular HA mRNA-LNP vaccines (black squares with dashed lines) or an 8-HA multivalent that contains all eight HA antigens (dark green circles). Titers were measured prior to vaccination at arrival, 14 days post vaccination (DPV), 14 days post boost (DPB) and 21 DPB. For challenged piglets, 21 DPB corresponds to the day of challenge (0 days post inoculation). (GraphPad Prism, GraphPad Software, La Jolla, CA).

**Table 1.**
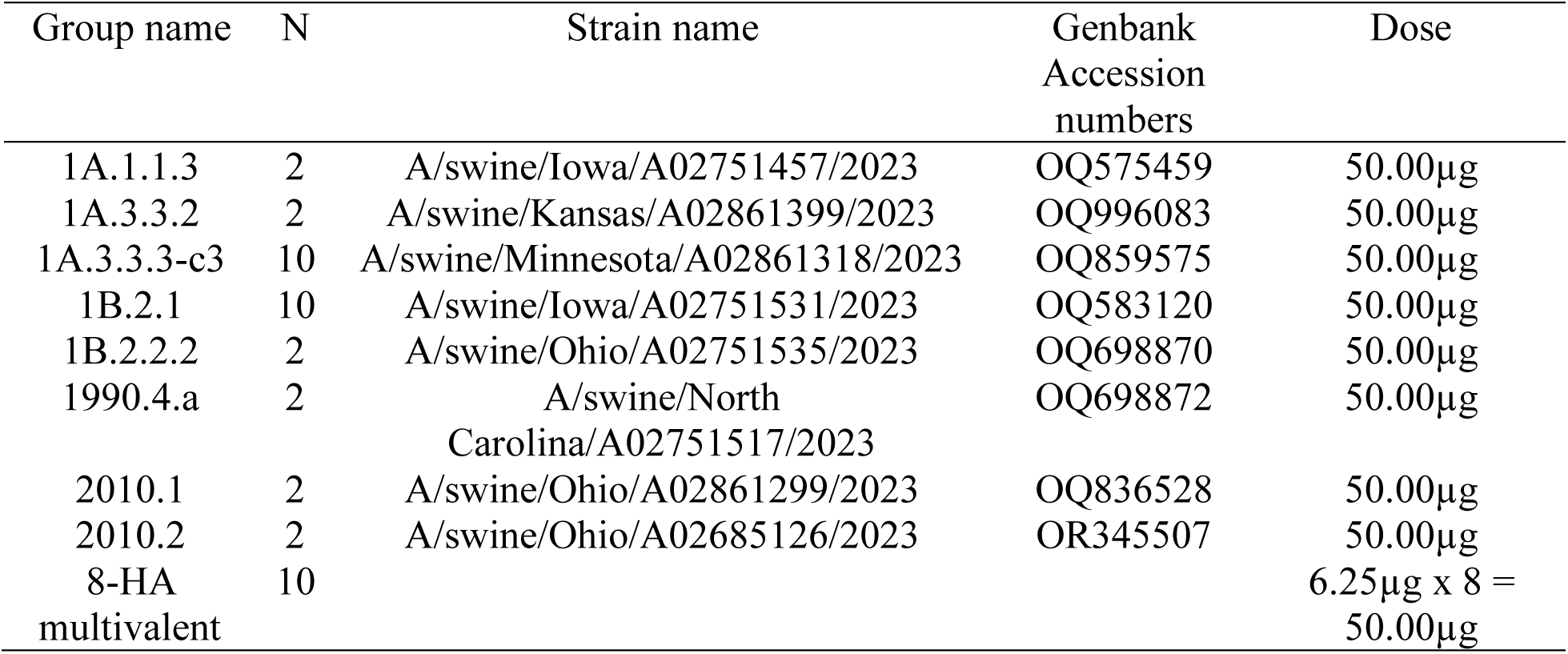
Vaccine target immunogenicity experimental group designations.

**Table 2.** Antigenic analysis: Swine H1 1A and 1B lineages.

|  |  | A/swine/Iowa/A02751457/2023 | A/swine/Kansas/A02861399/2023 | A/swine/Minnesota/A02861318/2023 | A/swine/Iowa/A02751531/2023 | A/swine/Ohio/A02751535/2023 | 8-HA Multivalent |
| --- | --- | --- | --- | --- | --- | --- | --- |
|  | Global Clade |  |  |  |  |  |  |
| A/swine/Iowa/A02751457/2023 | 1A.1.1.3 | 1280 | <10 | <10 | 10 | 10 | 53 |
| A/swine/Kansas/A02861399/2023 | 1A.3.3.2 | <10 | 5120 | 40 | <10 | <10 | 106 |
| A/swine/Minnesota/A02861318/2023 | 1A.3.3.3-c3 | <10 | 14 | 1810 | <10 | 20 | 28 |
| A/swine/Iowa/A02751531/2023 | 1B.2.1 | <10 | <10 | <10 | 1810 | 20 | 56 |
| A/swine/Ohio/A02751535/2023 | 1B.2.2.2 | <10 | <10 | <10 | <10 | 640 | 43 |

Hemagglutination inhibition (HI) from singular HA or 8-HA multivalent immunized swine antisera (columns) against endemic swine H1 1A and 1B IAV lineages (rows). Values are shown as geometric mean titers (GMT) at 21 days post-boost. Grey boxes indicate homologous GMT while red boxes indicate greater than eight-fold reduction in HI titers to the homologous value.

**Table 3.** Antigenic analysis : Swine H3s 1990.4, 2010.1 and 2010.2 lineages.

|  |  | A/swine/North<br>Carolina/A02751517/2023 | A/swine/North<br>Carolina/A02751517/2023 | A/swine/North<br>Carolina/A02751517/2023 | 8-HA Multivalent |
| --- | --- | --- | --- | --- | --- |
|  | Global<br>Clade |  |  |  |  |
| A/swine/North Carolina/A02751517/2023 | 1990.4.a | 905 | 40 | 10 | 70 |
| A/swine/Ohio/A02861299/2023 | 2010.1 | 28 | 1810 | 56 | 74 |
| A/swine/Ohio/A02685126/2023 | 2010.2 | 10 | 113 | 7241 | 226 |

Hemagglutination inhibition (HI) from singular HA or 8-HA multivalent immunized swine antisera (columns) against endemic swine H3 1990.4a, 2010.1 and 2010.2 IAV lineages (rows). Values are shown as geometric mean titers (GMT) at 21 days post-boost. Grey boxes indicate homologous GMT while red boxes indicate greater than eight-fold loss in HI titers to the homologous value.

### Piglets had robust HI antibody titers when vaccinated with low and high dose monovalent mRNA-LNP vaccines

For the challenge portion of the study, IAV naïve piglets were vaccinated and boosted intramuscularly with either a 6.25 µg dose (N=5) or a 50 µg dose (N=10) of monovalent HA mRNA-LNP vaccines to compare with the 8-HA multivalent vaccine recipients that received a total dose of 50 µg by including 6.25 µg dose of each individual mRNA-LNP. The monovalent vaccine components were formulated to encode the 1A.3.3.3-c3 HA clade representative homologous to the challenge strain (A/swine/Minnesota/A02861318/2023) or the 1B.2.1 HA clade representative (A/swine/Iowa/A02751531/2023) heterologous to the challenge strain. Serum was collected to evaluate antibody titers and cross-reactivity by HI assays (Figure 3). All vaccine recipients, from both low- and high-dose monovalent groups, had HI antibody titers greater than 1:40 against their respective homologous HA clade representative virus at 14 days post-boost (DPB) and maintained at the day of challenge (0 DPI). As expected, antibodies elicited by the 1A.3.3.3-c3 monovalent mRNA-LNP vaccine had limited HI antibody cross-reactivity to the 1B.2.1 test antigen and vice versa, regardless of the dose.

**Figure 3.**
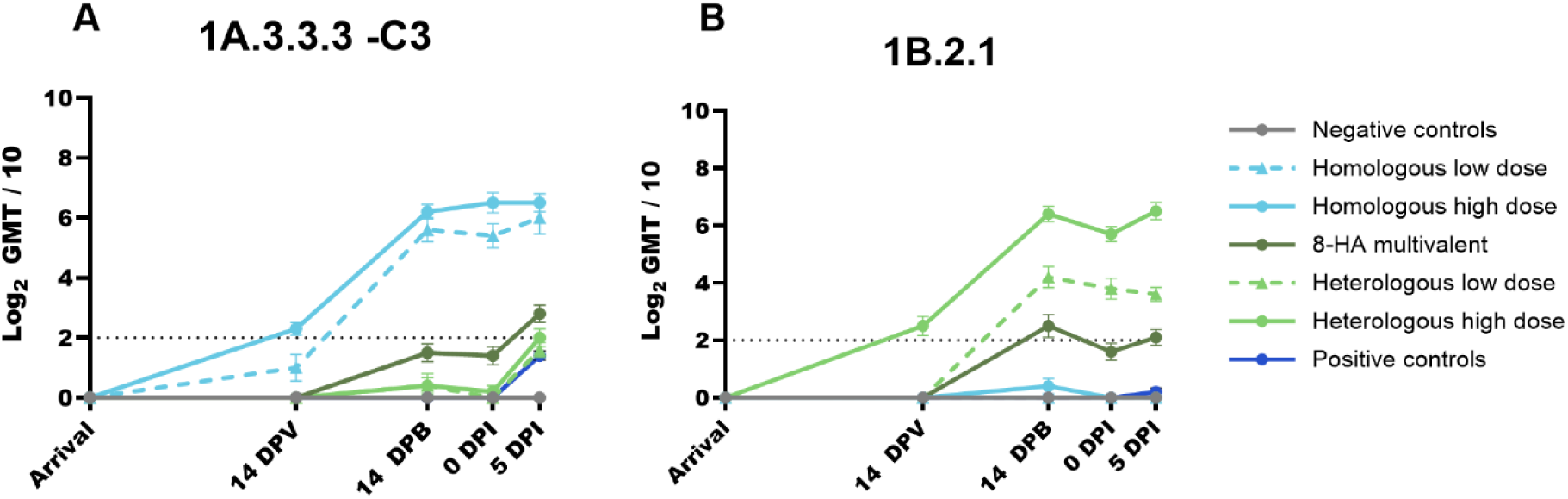
Hemagglutination inhibition (HI) antibody responses against 1A.3.3.3-c3 (A) and 1B.2.1 (B) strains. Log_2_-transformed mean HI titers are shown for pigs vaccinated with homologous (light blue) or heterologous (light green) to the challenge strain in low- (triangle) and high-dose (circle) as singular HA mRNA-LNP vaccines or the 8-HA multivalent (dark green) that contains all eight antigens. Negative control (grey) and positive control (dark blue) piglets that did not receive vaccines are included. Titers were measured at prior to vaccination at arrival, 14 days post vaccination (DPV), 14 days post boost (DPB), at challenge (0 DPI) and at necropsy (5 DPI). 0 DPI corresponds to 21 DPB. (GraphPad Prism, GraphPad Software, La Jolla, CA).

### Homologous mRNA–LNP vaccination significantly reduced virus detection in nasal secretions and the lower respiratory tract, independent of amount of mRNA

Piglets that received either the low-dose or the high-dose monovalent 1A.3.3.3-c3 and 1B.2.1 mRNA-LNP vaccines, as well as those that received the 8-HA multivalent mRNA-LNP vaccine, were subjected to an intranasal and intratracheal inoculation of 1A.3.3.3-c3 IAV at 21 DPB. For comparison, positive controls (non-vaccinated and challenged, N=10) were also inoculated. Nasal swabs were collected daily for 5 days post-inoculation (DPI) to detect shedding. Bronchioalveolar lavage fluid (BALF) was collected at necropsy to evaluate IAV detection in the trachea and lung. Virus titers for nasal swabs and BALF samples are presented in Figure 4; RT-PCR Ct values presented in Supplemental Figure 1. Negative control piglets remained IAV antibody negative throughout the study and did not have positive detections of IAV in their nasal swabs or BALF. The low- and high-dose homologous mRNA-LNP vaccines delayed the onset of nasal shedding and significantly reduced nasal viral titers at all DPI compared with the positive controls and heterologous vaccine groups. Piglets administered the 8-HA multivalent vaccine demonstrated a trend toward reduced IAV detection in nasal secretions, with titers generally lower than those of heterologous vaccine recipients and positive controls during peak shedding. Piglets that received either the low- or the high-dose heterologous vaccine had the highest and most sustained nasal viral titers, with group means statistically similar to the positive control group at all DPI. Mean cumulative Log_10_ TCID_50_/ml nasal viral titers across 1-5 DPI were lowest in the high-dose homologous group (0.52), followed by the low-dose homologous group (1.23), the 8-HA multivalent mRNA-LNP (2.69), and the positive controls (3.16), and were highest in low-dose and high-dose heterologous challenged recipients (3.58 and 3.47).

**Figure 4.**
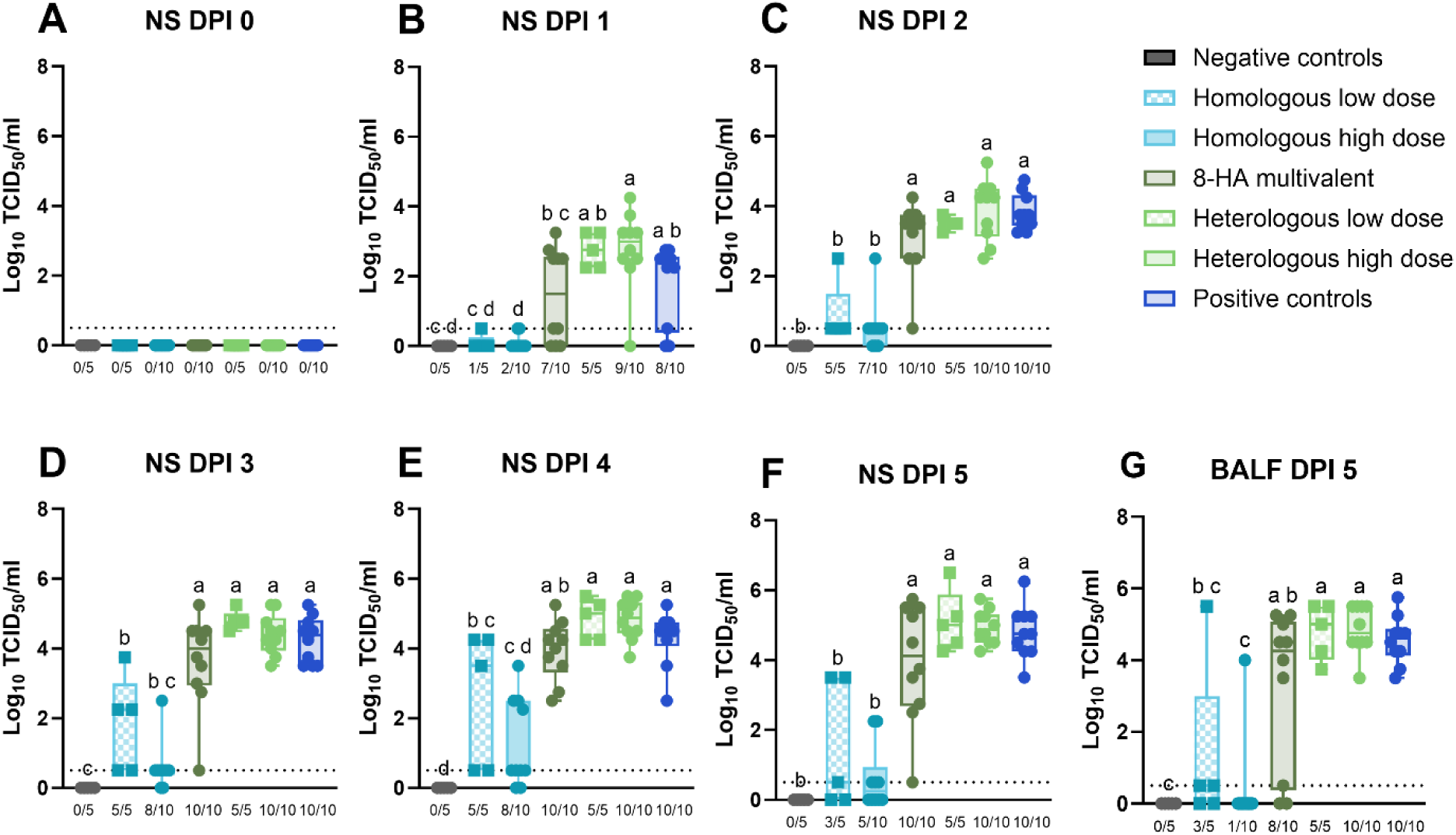
Virus titration on MDCK cells. Influenza A virus (IAV) titers from nasal swabs (A-F) collected 0-5 days post-inoculation (DPI) and BALF (G) collected at 5 DPI. Virus titration results are reported as log_10_ TCID_50_/ml in box and whisker plots showing all points. Numbers underneath the plots show positive pigs out of the group total. Different lower-case letters (a,b,c) indicate statistically significant difference (p ≤ 0.05) by ordinary one-way ANOVA with Tukey’s multiple comparisons test (GraphPad Prism, GraphPad Software, La Jolla, CA).

All BALF samples were positive for IAV qRT-PCR in the heterologous low- and high-dose groups and the positive controls (Supplemental data file). There was a significant difference between the group mean Log_10_ TCID_50_/ml virus titers isolated from the lungs of piglets that received the homologous low-dose (1.3) and homologous high-dose (0.4) vaccines in comparison to those that received the heterologous low-dose (4.8) and high-dose (4.9) vaccines, and to the positive control piglets (4.6). Piglets that received the 8-HA multivalent mRNA-LNP vaccine had intermediate levels of virus detected in the lungs (group average of 3.3 Log_10_ TCID_50_/ml), and this value was statistically similar to all challenge viral titers in BALF except the high-dose homologous recipients (p= 0.0005).

### Homologous and 8-HA mRNA-LNP vaccination protected piglets from lung lesions, whereas heterologous vaccination lacked cross-protection but did not induce VAERD

After inoculation, piglets were monitored for clinical signs of respiratory disease, including fever, coughing, anorexia, lethargy, respiratory distress, and eye irritation characterized by conjunctivitis and ocular discharge (Supplemental data file). At 3-5 DPI, three pigs in the high-dose heterologous group and one pig in the positive control group at 4 DPI exhibited respiratory distress after handling for sample collection; otherwise, no clinical signs were reported in the remaining vaccine treatment groups after inoculation. At necropsy, macroscopic evaluation of individual lungs was performed to score lesions with purple-red areas of consolidation typical of IAV. Piglets vaccinated with the 8-HA multivalent mRNA-LNP vaccine, as well as the low-dose and high-dose homologous monovalent vaccines, exhibited statistically significant lower lung lesion scores than the non-vaccinated, positive control animals (Figure 5). Low-dose and high-dose heterologous vaccine recipients had intermediate scores for macroscopic lung lesions, which were statistically similar to those of all groups except the non-challenged negative control. The weighted average percentages of macroscopic lung lesions were the lowest in the homologous vaccine low- and high-dose groups (0.1% for both dosages, similar to the negative controls), followed by 8-HA multivalent mRNA-LNP (0.4%), heterologous high-dose (2.0%), heterologous low-dose (2.6%), and positive controls (3.1%).

**Figure 5.**
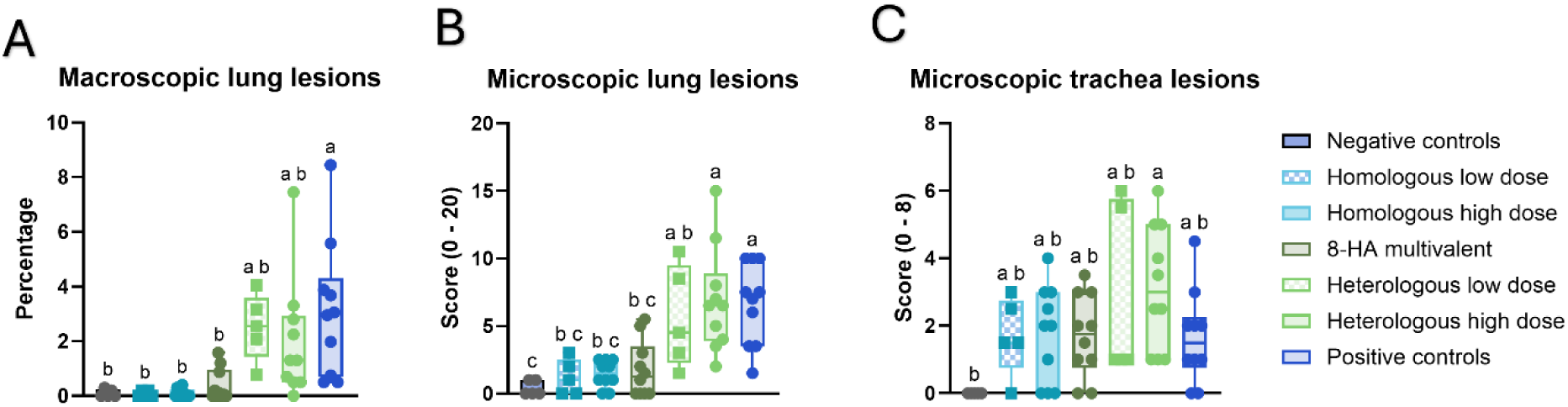
Respiratory tract lesion scores. (A) At necropsy, the percentage of lung surface area containing lesions typical of IAV infection was recorded and transformed into weighted percentages of macroscopic lung lesions. (B) Microscopic lung lesions (0-20) based on severity of interstitial pneumonia, peribronchiolar lymphocytic cuffing, bronchiolar epithelial necrosis, suppurative bronchitis and bronchiolitis, exocytosis and edema. (C) Microscopic trachea lesions scores (0-8) based on severity of tracheal epithelial necrosis and submucosal inflammation. Different lower-case letters (a,b,c) indicate statistically significant difference (p ≤ 0.05) by ordinary one-way ANOVA with Tukey’s multiple comparisons test (GraphPad Prism, GraphPad Software, La Jolla, CA).

Histopathological analysis of the lungs demonstrated a similar pattern, and piglets in the 8-HA multivalent mRNA-LNP, low-dose, and high-dose homologous monovalent mRNA-LNP vaccine groups had significantly lower scores for microscopic lung lesions than those in the heterologous monovalent vaccines and non-vaccinated, positive control groups. The highest group mean microscopic lung lesion scores were from high-dose heterologous group (6.9 ± 1.23), followed by the positive control group (6.7 ± 1.0), low-dose heterologous (5.6 ± 1.7) compared to the 8-HA multivalent mRNA-LNP (1.8 ± 0.7), high -dose homologous mRNA-LNP (1.5 ± 0.6), and low-dose homologous mRNA-LNP (1.2 ± 0.6), and negative controls (0.4 ± 0.2). All IAV inoculated groups, independent of vaccine, had mild to moderate tracheal epithelial necrosis and tracheitis histopathological scores, and there were no statistically significant differences between the reported group means.

## Discussion

Controlling IAV in North American swine through vaccination remains a challenge due to frequent reassortment, rapid antigenic drift, and prolonged co-circulation of genetically distinct H1 and H3 lineages and groups. Here, we evaluated a novel approach to enhance antigenic coverage against relevant strains of swine IAV. We designed a multivalent HA mRNA-LNP-based influenza vaccine that maximized coverage of genetic diversity through a phylogenetically informed selection of representative strains from genomic surveillance data. We demonstrated that the simultaneous delivery of eight distinct IAV antigens generated HA-specific antibodies that react to over 90% of the current observed swine HA genetic diversity in the U.S. We also demonstrated that this vaccine platform protected piglets against IAV clinical disease and lung lesions when challenged with a wild type homologous strain, selected as it is representative of 20% of H1s currently detected in North American swine. The recipients of the 8-HA multivalent mRNA-LNP vaccine had average macroscopic and microscopic lung lesions statistically similar to the negative control piglets and homologous vaccine recipients.

Although the 8-HA multivalent pigs had detectable HI antibody titers, only the piglets that received the homologous monovalent formulations of the mRNA-LNP vaccine had a significant reduction of IAV in nasal secretions. There is a possibility that to the lack of protection against nasal viral shedding was due to amount of antigen delivered per each HA in the multivalent vaccine formulation. Therefore, there is a need to understand the optimal number of antigens and individual dosages that can be simultaneously delivered through mRNA-LNPs as well as the overall dose of a multivalent mRNA-LNP vaccine that will result in HI titers with efficacy against virus replication in the upper respiratory tract. Intramuscular delivery of the mRNA-LNP vaccines might explain their limited impact on viral replication in the nasal mucosa. In general, intramuscular vaccination stimulates systemic IgM and IgG antibodies that circulate in serum and can transudate into the lung during the acute phase of infection, but this transudation is less effective in the mucosa of the upper respiratory tract, including nasal turbinates and trachea [8]. Intramuscular vaccination does not stimulate strong mucosal IgA responses. Although cellular immune responses and mucosal IgA levels were not evaluated in the present study, they may contribute to reduced shedding and are known to exhibit greater cross-reactivity [24, 25]. Previous studies in the swine model have shown that intranasal immunization with LNP-encapsulated plasmid DNA encoding HA proteins induced stronger mucosal IgA and cellular responses than intramuscular immunization, while also reducing nasal viral shedding [26]. Therefore, future studies should investigate mucosal delivery methods for multivalent mRNA-LNP vaccines or strategies to enhance mucosal and cellular immunity and reduce shedding. These strategies could include adding immunomodulators to the vaccine formulation. Previous research with live attenuated IAV (LAIV) vaccines in swine demonstrated that the addition of IgA-inducing protein (IGIP) to the vaccines reduced nasal viral shedding and completely blocked transmission to naïve contacts compared with a non-IGIP LAIV formulation [25]. In addition, modifications of mRNA-LNP formulations to include cytokines, such as IL-12, have been shown to enhance CD8 T cell-mediated responses against IAV and represent another strategy to shape the magnitude of the response to vaccination [27].

The implications of sequential vaccination and challenge with mismatched IAV between the H1 1A and 1B lineages is a concern in pigs [12]. We included a heterologous vaccination group in the study to evaluate whether mRNA-LNP vaccination would induce VAERD after challenge with a mismatched IAV strain. The VAERD phenomenon occurs when a pig is vaccinated with an adjuvanted WIV and is infected with an antigenically distinct strain of the same subtype. Cross-reactive antibodies that lack neutralizing activity, then activate the complement system and dysregulate proinflammatory cytokines, thereby actually increasing the severity of clinical disease and lung lesions [14]. In this study, the mRNA-LNP mismatched vaccine did not induce VAERD. Piglets in the 1B.2.1 heterologous vaccine groups had macroscopic and microscopic lung lesions that were not elevated above those of the non-vaccinated (positive control) piglets and did not demonstrate exacerbated clinical disease scores after being infected with an 1A.3.3.3-c3 strain. Alphavirus-derived RP vaccines containing influenza HA RNA also provides similar benefits to the mRNA-LNP platform including robust protection without inducing VAERD [17] and a future study that directly compares immunogenicity and efficacy between these two nucleic acid platforms would be of interest.

Our results emphasize the benefits of mRNA-LNP vaccines for controlling IAV in swine. We demonstrated that these vaccines were safe, immunogenic, effective at protecting the piglets against challenge, and did not induce VAERD. The 8-HA multivalent mRNA-LNP vaccination induced an HI antibody response to all 8 HA targets that was boosted to the challenge strain after challenge, suggesting that priming piglets with a breadth of HAs will better prepare them against subsequent infections with a diversity of IAV by shortening the time to production of protective antibodies. In addition, the mRNA vaccine platform has several advantages over other experimentally tested IAV vaccines, such as subunit, WIV, LAIV, and DNA-based vaccines [15, 16]. Unlike WIV, once the HA sequences of circulating influenza virus strains are known, mRNA-LNP vaccines can be easily updated without the complications of cell or egg culture. The LNP supplies potent adjuvant activity that selectively stimulates a T helper 1-biased T follicular helper cell response [28, 29]. Unlike plasmid DNA, mRNA cannot integrate into the host genome and poses no danger of genomic toxicity. In addition, mRNA is short-lived, unlike recombinant viruses, limiting vector persistence and antigen expression; therefore, it allows for precise control over pharmacokinetics and dosing [15, 30]. Another benefit of the vaccine platform is that recipients of HA or NA sequence-based vaccines can be serologically distinguished from naturally infected individuals through antibody tests targeting the viral NP, which is commonly used for IAV serodiagnosis [13].

Although the results are encouraging and highlight advantages of this novel vaccine approach, several considerations and aspects regarding the use of mRNA-LNP vaccines to control swine IAV remain unaddressed. While HA-focused immunity is critical for virus neutralization, the incorporation of additional antigens such as neuraminidase (NA) or conserved internal proteins may further enhance cross-protection and expand the length to which these vaccines might be relevant to use in the field [31]. Also, the long-term durability of vaccine-induced protection was not evaluated in this study, as piglets were euthanized at 26 days post-boost. In the swine industry, gilts are often vaccinated against IAV twice before entering the breeding herd, and sows are vaccinated quarterly before farrowing to pass MDA to their piglets [32]. These production practices raise three important considerations for the implementation of swine mRNA-LNP vaccines in commercial swine. First, the optimal prime-boost strategy remains to be determined. In our study, we used the same formulation and delivery method to prime and boost vaccinated piglets. However, previous research has shown that heterologous prime-boost approaches enhance the magnitude and breadth of IAV immune responses compared to homologous boost strategies [33, 34]. Therefore, exploration of heterologous prime-and-boost strategies may broaden protection against IAV and allow for reducing the number of HA antigens per dose without sacrificing cross-reactivity [19]. Second, a sow is considerably larger than a piglet; therefore, an investigation into the dosage of mRNA needed to induce appropriate immune responses in older, larger animals is warranted. Third, the impact of MDA on mRNA vaccine performance needs further investigation, particularly given the challenges MDA poses for conventional WIV HA-based vaccines in young piglets [35]. Studies using an LNP-encapsulated plasmid DNA vaccine expressing HA demonstrated that MDA did not significantly interfere with vaccine-induced responses or protection in piglets, which may suggest a similar outcome for mRNA-LNP vaccines [13]. In addition, previous research done in mouse pups demonstrated that mRNA-LNP vaccines elicited protective antibodies against IAV in the presence of maternal antibodies [36].

Collectively, our data support the use of monovalent and multivalent mRNA-LNP vaccines targeted against the HA of IAV in swine, informed by production systems or regional surveillance, as a promising, quickly adaptable strategy for inducing robust and broad immunity in swine. The application of these vaccines may lead to superior vaccine programs that improve animal health, minimize the diversity of endemic IAV and reduce the risk to humans by reducing the viral burden in swine populations, as well as reassortment and the emergence of novel strains with zoonotic potential.

## Materials and Methods

### Hemagglutinin gene selection and quantifying representation of genetic diversity

Strain metadata and HA nucleotide sequence data collected in the U.S. from January 2022 to April 2025 were queried from GenBank and GISAID databases. The dataset was cleaned, removing duplicates and erroneous sequences, and split by HA subtype then cropped to the HA1 subunit before aligning with MAFFT v7.526 and building phylogenetic trees with FastTree v2.1.4. For each monophyletic group of viruses identified on the inferred phylogeny, a single representative selection was made using PARNAS v0.1.7 [23]. We selected 8 antigens from 14 potential HA groups that were identified within the USDA IAV in swine surveillance system between 2022 and 2025, as these 8 clades represented 95.8% of detections [6, 37]. Subsequently, we used PARNAS v0.1.7 to rescale the branches of the H1 and H3 HA trees to infer ancestral HA1 amino acid substitutions along each branch, allowing coverage for each of the selected HA vaccine antigens to be determined at a 16 amino acid (∼5%) difference threshold; this threshold was used as a correlate of antigenic similarity [5].

### mRNA-LNP production and lipid nanoparticle (LNP) encapsulation

HA gene segments from eight genetic clades of swine-lineage IAV were utilized for antigen design, as previously described [22, 38, 39]. Briefly, HA sequences were codon-optimized for improved mRNA translation and half-life [40] and cloned into mRNA production vectors. The HA coding sequence (CDS) is flanked by a T7 promoter, a 5’-untranslated region (UTR) derived from the tobacco etch virus (TEV), and a Kozak sequence upstream cassette and a 3’-UTR of Xenopus beta-globin and a 101-nucleotide long poly(A) tail downstream cassette [41]. Vectors were linearized and mRNAs were produced using a T7-driven in vitro transcription reaction (Megascript, Ambion). m1Ψ-5′-triphosphate (TriLink) was incorporated into the reaction instead of UTP. In vitro transcribed mRNAs were co-transcriptionally capped using CleanCap (TriLink) [38] and purified by a fast protein liquid chromatography (FPLC)-based cellulose chromatography [42]. All purified mRNAs were analyzed using native agarose gel electrophoresis, tested for dsRNA and endotoxin content using dot blot and LAL chromogenic assay, respectively, and stored at −20°C. Purified mRNAs were encapsulated in LNP using a self-assembly process, as previously described [43], in which an aqueous mRNA-containing solution at acidic pH is rapidly mixed with an ethanolic lipid mixture of ionizable cationic lipid (proprietary to Acuitas), phosphatidylcholine, cholesterol, and polyethylene glycol (PEG)–lipid (50:10:38.5:1.5 mol/mol). mRNA-LNP formulations were subsequently stored at −80°C until used. The mean hydrodynamic diameter of these mRNA-LNPs was ∼80 nm an encapsulation efficiency of ∼95% (Acuitas).

### Viruses

The eight IAV isolates selected as clade representatives originated from diagnostic investigations of clinical cases of respiratory disease in pigs and were obtained from the repository at the National Veterinary Services Laboratories (NVSL) through the U.S. Department of Agriculture (USDA) National Surveillance Plan for Swine Influenza Virus [37]. The selected viruses covered the H1 1A classical swine lineage, A/swine/Iowa/A02751457/2023 (1A.1.1.3), A/swine/Kansas/A02861399/2023 (1A.3.3.2), A/swine/Minnesota/A02861318/2023 (1A.3.3.3-c3); the H1 1B human seasonal lineage, A/swine/Iowa/A02751531/2023 (1B.2.1), A/swine/Ohio/A02751535/2023 (1B.2.2.2); and the three major H3 subtype lineages detected in pigs in the U.S., A/swine/North Carolina/A02751517/2023 (1990.4.a), A/swine/Ohio/A02861299/2023 (2010.1) and A/swine/Ohio/A02685126/2023 (2010.2). All isolates were propagated and titrated in Madin-Darby canine kidney (MDCK) cells using Opti-MEM™ (Life Technologies, Waltham, MA) containing antibiotics/antimycotics and 1µg/ml of tosyl sulfonyl phenylalanyl chloromethyl ketone (TPCK)-trypsin (Worthington Biochemical Corp., Lakewood, NJ) infection media.

### Vaccination study design

Sixty-seven three-week-old pigs were obtained from a high-health commercial source free of porcine reproductive and respiratory syndrome virus (PRRSV), Mycoplasma hyopneumoniae (MHP), and IAV. After arrival, they were housed in biosafety level 2 containment and cared for in compliance with the Institutional Animal Care and Use Committee of Iowa State University (IACUC # 25-145 and 25-146). All animals were treated with enrofloxacin (Elanco, Indianapolis, IN) to decrease the risk of pre-existing bacterial infections and were screened by a commercial NP ELISA kit (Swine Influenza Virus Antibody Test, IDEXX, Westbrook, ME) to ensure the absence of IAV antibodies. Pigs were implanted with a Life Chip from Bio-Thermo Technology (Destron Fearing, DFW Airport, Texas). Nasal swabs were collected three days post-arrival to confirm the IAV negative status by qRT-PCR. At 4 weeks of age, pigs were randomly divided into thirteen vaccination treatment groups as shown in Table 1. The vaccine protocol involved two doses of mRNA-LNP vaccine with an interval of three weeks between prime and boost. mRNA-LNPs (6.25 µg for low-dose or 50 µg for high-dose) were diluted into 1 ml of sterile PBS and injected intramuscularly into the right side of the neck. For the 8 HA mRNA-LNP vaccinations, individual HA mRNA-LNPs were pooled before vaccination (comprised of 6.25 µg of each individual HA combined into a dose of 50 µg). Blood samples were collected at 0, 21, 35, and 42 DPV to evaluate serological immune responses.

### Challenge study design

Two weeks post-boost, piglets were inoculated with 2ml intratracheally and 1ml intranasally with 1 × 10^5^ 50% tissue culture infectious dose (TCID_50_)/ml of 1A.3.3.3-c3 clade representative strain (A/swine/Minnesota/A02861318/2023) as shown in Table 4. Inoculation was done under sedation, using an intramuscular injection of ketamine (2.2 mg/kg of body weight; Phoenix, St. Joseph, MO), xylazine (2.2 mg/kg; Lloyd Inc., Shenandoah, IA), and Telazol (4.4 mg/kg; Zoetis Animal Health, Florham Park, NJ) cocktail. Nasal swabs (NS) were collected daily from 0 to 5 DPI in media containing 2 ml minimum-essential-medium (MEM). On 5 DPI, pigs were bled and humanely euthanized with a lethal dose of pentobarbital (Fatal Plus; Vortech Pharmaceuticals, Dearborn, MI). Lungs were aseptically removed, evaluated for macroscopic lesions, and lavaged with 50 ml of MEM to obtain BALF. A section of the right middle or affected lung lobe and distal trachea were collected and fixed in 10% buffered formalin for microscopic examination and scoring.

**Table 4.** Vaccine efficacy experimental group designations.

| Group name | N | mRNA-LNP vaccine target | Dose | Challenge strain |
| --- | --- | --- | --- | --- |
| Negative controls | 5 | N/A | N/A | N/A |
| Homologous low dose | 5 | 1A.3.3.3-c3 | 6.25µg | 1A.3.3.3-c3 |
| Homologous high dose | 10 | 1A.3.3.3-c3 | 50.00µg | 1A.3.3.3-c3 |
| Heterologous low dose | 5 | 1B.2.1 | 6.25µg | 1A.3.3.3-c3 |
| Heterologous high dose | 10 | 1B.2.1 | 50.00µg | 1A.3.3.3-c3 |
| 8-HA multivalent | 10 | 1A.1.1.3, 1A.3.3.2, 1A.3.3.3-c3, 1B.2.1, 1B.2.2.2, 1990.4.a, 2010.1, 2010.2. | 6.25µg x 8 = 50.00µg | 1A.3.3.3-c3 |
| Positive controls | 10 | N/A | N/A | 1A.3.3.3-c3 |

### Hemagglutination inhibition assays

Serum samples were obtained from the pigs at 0, 21, 35, and 42 days DPV and 5 DPI to test for neutralizing HA antibodies by HI assay. Prior to HI, sera were heat inactivated at 56℃ for 30 minutes, incubated with 20% kaolin in PBS solution for 20 minutes, and adsorbed with 0.5% and subsequently 100% turkey red blood cells for 20 minutes each to remove nonspecific hemagglutination inhibitors and natural serum agglutinins. Homologous and heterologous HI assays were performed using standard techniques [10]. The reciprocal titers were divided by 10, log_2_ transformed, and reported as the geometric mean.

### Virus detection

All nasal swabs and BALF samples were submitted to the Iowa State University Veterinary Diagnostic Laboratory (ISU VDL) for IAV qRT-PCR testing. Samples with cycle threshold (Ct) values below 38 were considered positive and subjected to virus isolation on 48-well and titration on 96-well plates of confluent MDCK cells. For virus isolation, cells were inoculated with 100µl of undiluted sample supplemented with TPCK-trypsin (1:1000). For virus titration, 10-fold serial triplicate dilutions of the sample were prepared in infection media, and 100µl of each dilution was inoculated in the subsequent rows. Plates were incubated at 37℃ for 48h hours, then fixed and stained for immunocytochemistry (ICC) staining as previously described [44]. Titers were calculated for each sample as TCID_50_/ml and transformed to log_10_ for comparison purposes. Samples that were positive in the virus isolation plate, but negative in the initial dilution of the virus titration received a score of 0.5.

### Lower respiratory tract lesion scoring

At necropsy, the percentage of lung surface area containing lesions typical of IAV infection was recorded. A visual estimate was made for each lung lobe, and a total percentage for the entire lung was calculated based on the weighted proportions of each lobe relative to the total lung volume [14, 45]. Lung and trachea tissues were submitted for routine histologic procedures at the ISU VDL, and the slides were stained with hematoxylin and eosin per standard operating procedures. Microscopic lesions were evaluated and scored by a veterinary pathologist blinded to group, using parameters previously described [14].

### Statistical analysis

Macroscopic and microscopic pneumonia scores, HI titer log_2_ geometric means, body temperatures, qRT-PCR Ct values, log_10_ transformed nasal swab and BALF viral titers were analyzed using analysis of variance (ordinary one-way ANOVA) with Tukey’s post-hoc multiple comparisons test of parameters with statistical differences. Mean cumulative Log_10_ TCID_50_/ml nasal viral titers across 1-5 DPI were calculated by summing the individual daily titers and dividing by 5. Comparisons were made between the groups at the same time point using a 5% significance level (p-value <0.05) to indicate statistically significant differences in GraphPad Prism (GraphPad Software, La Jolla, CA).

### Data availability statement

Clinical data associated with this study are available for download from the USDA Ag Data Commons: DOI: X. All genetic sequence data are available at NCBI GenBank with USDA IAV in swine surveillance data within a searchable interface at https://flu-crew.org.

## Acknowledgments

We thank Emily Love, Katharine Young, Emily Church and Alissa Hufnagel for their laboratory assistance; and Marcelo Almeida, Jianqiang Zhang, Joe Thomas, Hu Ke and Iowa State University Laboratory Animal Resources caretaker staff for their support with the animals. We gratefully acknowledge pork producers, swine veterinarians, and laboratories for participating in the USDA Influenza A Virus in Swine Surveillance System. Funding was provided in part by NIH NIAID CEIRR (#75N93021C00015) and USDA-ARS (ARS project number 5030-32000-231-000D). This research was supported in part by an appointment to the ARS Research Participation Program administered by the Oak Ridge Institute for Science and Education (ORISE) through an interagency agreement between the U.S. Department of Energy (DOE) and the USDA. ORISE is managed by ORAU under DOE contract number DE-SC0014664. All opinions expressed in this paper are the author’s and do not necessarily reflect the policies and views of USDA, DOE, or ORAU/ORISE. The use of trade or corporation names in this publication is for information purposes and does not constitute an official endorsement by the USDA ARS to the exclusion of others. USDA is an equal opportunity provider and employer.

---

**Supplemental Figure 1.**
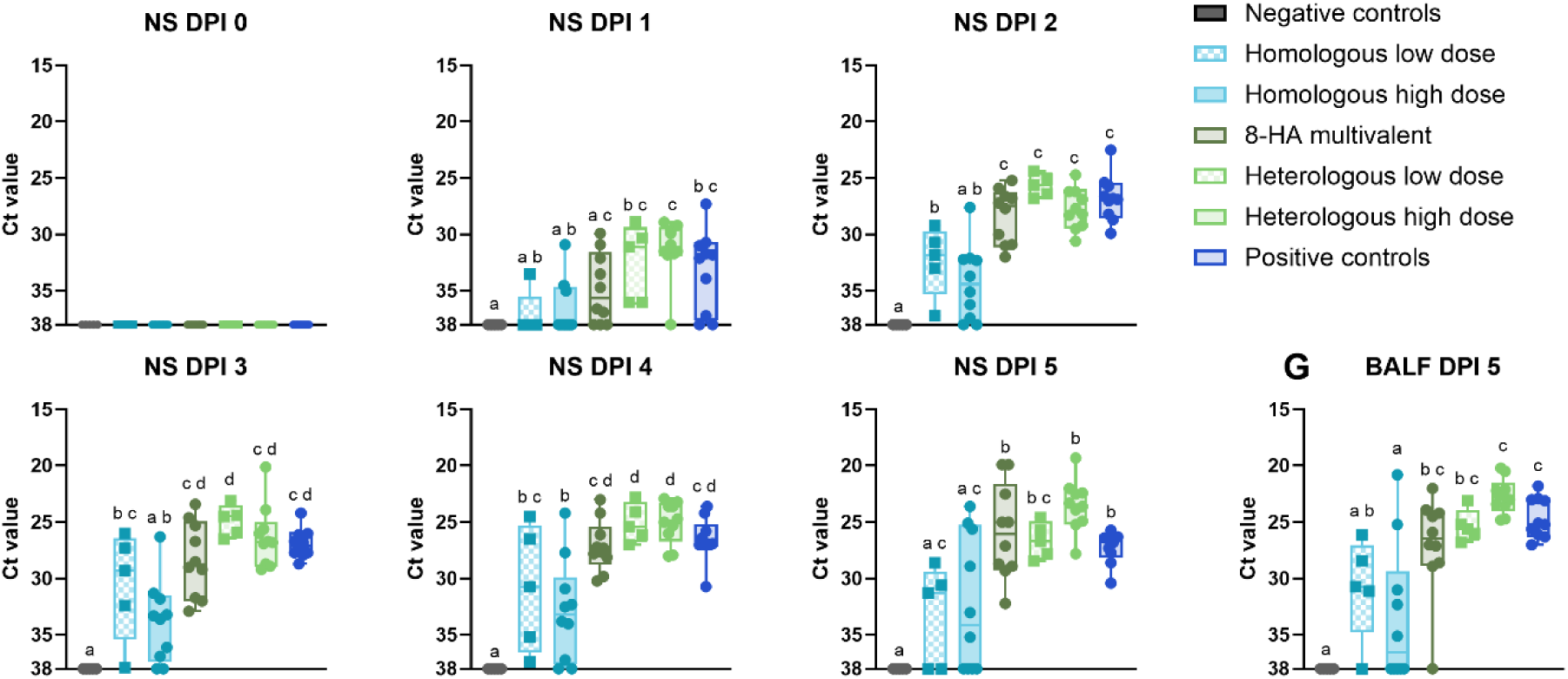
IAV detection by qRT-PCR on nasal swabs (A-F) collected 0-5 days post-inoculation (DPI) and BALF (G) collected at 5 DPI. qRT-PCR results are reported as Ct values in box and whisker plots showing all points. Different lower-case letters (a,b,c) indicate statistically significant difference (p ≤ 0.05) by ordinary one-way ANOVA with Tukey’s multiple comparisons test (GraphPad Prism, GraphPad Software, La Jolla, CA).

## Notes

### Competing Interest Statement

The authors have declared no competing interest.

## References

1. Vincent, A.L., et al., Swine influenza viruses a North American perspective. Adv Virus Res, 2008. 72: p. 127–54.

2. Steel, J. and A.C. Lowen, Influenza A virus reassortment. Curr Top Microbiol Immunol, 2014. 385: p. 377–401.

3. Zhou, N.N., et al., Genetic reassortment of avian, swine, and human influenza A viruses in American pigs. J Virol, 1999. 73(10): p. 8851–6.

4. Garten, R.J., et al., Antigenic and Genetic Characteristics of Swine-Origin 2009 A(H1N1) Influenza Viruses Circulating in Humans. Science, 2009. 325(5937): p. 197–201.

5. Anderson, T.K., et al., Swine Influenza A Viruses and the Tangled Relationship with Humans. Cold Spring Harb Perspect Med, 2021. 11(3).

6. Arendsee, Z.W., et al., octoFLUshow: an Interactive Tool Describing Spatial and Temporal Trends in the Genetic Diversity of Influenza A Virus in U.S. Swine. Microbiol Resour Announc, 2021. 10(50): p. e0108121.

7. Van Reeth, K. and A.L. Vincent, Influenza Viruses, in Diseases of Swine. 2019. p. 576–593.

8. Van Reeth, K. and W. Ma, Swine Influenza Virus Vaccines: To Change or Not to Change—That’s the Question, in Swine Influenza, J.A. Richt and R.J. Webby, Editors. 2013, Springer Berlin Heidelberg: Berlin, Heidelberg. p. 173–200.

9. Vincent, A.L., et al., Influenza A virus vaccines for swine. Veterinary Microbiology, 2017. 206: p. 35–44.

10. Kitikoon, P., P.C. Gauger, and A.L. Vincent, Hemagglutinin inhibition assay with swine sera. Methods Mol Biol, 2014. 1161: p. 295–301.

11. Thomas, M.N., et al., Active surveillance for influenza A virus in swine reveals within-farm reassortment and cocirculation of distinct subtypes and genetic clades. Veterinary Microbiology, 2025. 309: p. 110681.

12. Gauger, P.C., et al., Enhanced pneumonia and disease in pigs vaccinated with an inactivated human-like (δ-cluster) H1N2 vaccine and challenged with pandemic 2009 H1N1 influenza virus. Vaccine, 2011. 29(15): p. 2712–2719.

13. Lai Danh, C., et al., Lipid nanoparticle-encapsulated DNA vaccine induces balanced antibody and T-cell responses in pigs with maternally derived antibodies. Journal of Virology, 2025. 99(11): p. e01123–25.

14. Gauger, P.C., et al., Kinetics of lung lesion development and pro-inflammatory cytokine response in pigs with vaccine-associated enhanced respiratory disease induced by challenge with pandemic (2009) A/H1N1 influenza virus. Vet Pathol, 2012. 49(6): p. 900–12.

15. Pardi, N., et al., mRNA vaccines — a new era in vaccinology. Nature Reviews Drug Discovery, 2018. 17(4): p. 261–279.

16. Pardi, N., M.J. Hogan, and D. Weissman, Recent advances in mRNA vaccine technology. Current Opinion in Immunology, 2020. 65: p. 14–20.

17. Wymore Brand, M., et al., Bivalent hemagglutinin and neuraminidase influenza replicon particle vaccines protect pigs against influenza a virus without causing vaccine associated enhanced respiratory disease. Vaccine, 2022. 40(38): p. 5569–5578.

18. Vander Veen, R.L., et al., Safety, immunogenicity, and efficacy of an alphavirus replicon-based swine influenza virus hemagglutinin vaccine. Vaccine, 2012. 30(11): p. 1944–1950.

19. Wymore Brand, M., et al., Swine influenza A replicon particle and live attenuated influenza virus vaccines induce differential systemic and mucosal antibody and T cell responses. Frontiers in Veterinary Science, 2026. Volume 12 - 2025.

20. Kitikoon, P., et al., Quadrivalent neuraminidase RNA particle vaccine protects pigs against homologous and heterologous strains of swine influenza virus infection. Vaccine, 2023. 41(47): p. 6941–6951.

21. Moraes, D.C.A., et al., Veterinarian perceptions and practices in prevention and control of influenza virus in the Midwest United States swine farms. Front Vet Sci, 2023. 10: p. 1089132.

22. Arevalo, C.P., et al., A multivalent nucleoside-modified mRNA vaccine against all known influenza virus subtypes. Science, 2022. 378(6622): p. 899–904.

23. Markin, A., et al., PARNAS: Objectively Selecting the Most Representative Taxa on a Phylogeny. Systematic Biology, 2023. 72(5): p. 1052–1063.

24. Loving, C.L., et al., Heightened adaptive immune responses following vaccination with a temperature-sensitive, live-attenuated influenza virus compared to adjuvanted, whole-inactivated virus in pigs. Vaccine, 2012. 30(40): p. 5830–5838.

25. Rajao, D.S., et al., Live attenuated influenza A virus vaccine expressing an IgA-inducing protein protects pigs against replication and transmission. Frontiers in Virology, 2023. Volume 3 - 2023.

26. Dhakal, S., et al., Liposomal nanoparticle-based conserved peptide influenza vaccine and monosodium urate crystal adjuvant elicit protective immune response in pigs. Int J Nanomedicine, 2018. 13: p. 6699–6715.

27. Aunins, E.A., et al., An Il12 mRNA-LNP adjuvant enhances mRNA vaccine-induced CD8 T cell responses. Science Immunology, 2025. 10(108): p. eads1328.

28. Alameh, M.G., et al., Lipid nanoparticles enhance the efficacy of mRNA and protein subunit vaccines by inducing robust T follicular helper cell and humoral responses. Immunity, 2021. 54(12): p. 2877–2892.e7.

29. Pardi, N., et al., Nucleoside-modified mRNA vaccines induce potent T follicular helper and germinal center B cell responses. J Exp Med, 2018. 215(6): p. 1571–1588.

30. Petsch, B., et al., Protective efficacy of in vitro synthesized, specific mRNA vaccines against influenza A virus infection. Nature Biotechnology, 2012. 30(12): p. 1210–1216.

31. van de Ven, K., et al., A universal influenza mRNA vaccine candidate boosts T cell responses and reduces zoonotic influenza virus disease in ferrets. Science Advances, 2022. 8(50): p. eadc9937.

32. Garrido-Mantilla, J., et al., Reduction of Influenza A Virus Prevalence in Pigs at Weaning After Using Custom-Made Influenza Vaccines in the Breeding Herds of an Integrated Swine Farm System. Viruses, 2025. 17(2): p. 240.

33. Dong, C., et al., Enhancing cross-protection against influenza by heterologous sequential immunization with mRNA LNP and protein nanoparticle vaccines. Nature Communications, 2024. 15(1): p. 5800.

34. Bhatnagar, N., et al., Heterologous Prime-Boost Vaccination with Inactivated Influenza Viruses Induces More Effective Cross-Protection than Homologous Repeat Vaccination. Vaccines (Basel), 2023. 11(7).

35. Allerson, M., et al., The impact of maternally derived immunity on influenza A virus transmission in neonatal pig populations. Vaccine, 2013. 31(3): p. 500–5.

36. Willis, E., et al., Nucleoside-modified mRNA vaccination partially overcomes maternal antibody inhibition of de novo immune responses in mice. Science Translational Medicine, 2020. 12(525): p. eaav5701.

37. Janzen, G.M., et al., Sources and sinks of influenza A virus genomic diversity in swine from 2009 to 2022 in the United States. Journal of Virology, 2025. 99(9): p. e00541–25.

38. Pardi, N., et al., Nucleoside-modified mRNA immunization elicits influenza virus hemagglutinin stalk-specific antibodies. Nature Communications, 2018. 9(1): p. 3361.

39. Gouma, S., et al., Nucleoside-Modified mRNA-Based Influenza Vaccines Circumvent Problems Associated with H3N2 Vaccine Strain Egg Adaptation. J Virol, 2023. 97(1): p. e0172322.

40. Thess, A., et al., Sequence-engineered mRNA Without Chemical Nucleoside Modifications Enables an Effective Protein Therapy in Large Animals. Mol Ther, 2015. 23(9): p. 1456–64.

41. Pardi, N., et al., In vitro transcription of long RNA containing modified nucleosides. Methods Mol Biol, 2013. 969: p. 29–42.

42. Baiersdorfer, M., et al., A Facile Method for the Removal of dsRNA Contaminant from In Vitro-Transcribed mRNA. Mol Ther Nucleic Acids, 2019. 15: p. 26–35.

43. Maier, M.A., et al., Biodegradable lipids enabling rapidly eliminated lipid nanoparticles for systemic delivery of RNAi therapeutics. Mol Ther, 2013. 21(8): p. 1570–8.

44. Gauger, P.C. and A.L. Vincent, Serum virus neutralization assay for detection and quantitation of serum-neutralizing antibodies to influenza A virus in swine. Methods Mol Biol, 2014. 1161: p. 313–24.

45. Halbur, P.G., et al., Comparison of the pathogenicity of two US porcine reproductive and respiratory syndrome virus isolates with that of the Lelystad virus. Vet Pathol, 1995. 32(6): p. 648–60.

